# Structure of monomeric full-length ARC sheds light on molecular flexibility, protein interactions, and functional modalities

**DOI:** 10.1101/332015

**Authors:** Erik I. Hallin, Maria S. Eriksen, Sergei Baryshnikov, Oleksii Nikolaienko, Sverre Grødem, Tomohisa Hosokawa, Yasunori Hayashi, Clive R. Bramham, Petri Kursula

## Abstract

The activity-regulated cytoskeleton-associated protein (ARC) is critical for long-term synaptic plasticity and memory formation. Acting as a protein interaction hub, ARC regulates diverse signalling events in postsynaptic neurons. A protein interaction site is present in the ARC C-terminal domain (CTD), a bilobar structure homologous to the retroviral Gag capsid domain. However, knowledge of the 3-dimensional structure of full-length ARC is required to elucidate its molecular function. We purified recombinant monomeric full-length ARC and analyzed its structure using small-angle X-ray scattering and synchrotron radiation circular dichroism spectroscopy. In solution, monomeric ARC has a compact, closed structure, in which the oppositely charged N-terminal domain (NTD) and CTD are juxtaposed, and the flexible linker between them is not extended. The modelled structure of ARC is supported by intramolecular live-cell FRET imaging in rat hippocampal slices. Peptides from several postsynaptic proteins, including stargazin, bind to the N-lobe, but not to the C-lobe, of the bilobar CTD. This interaction does not induce large-scale conformational changes in the CTD or flanking unfolded regions. The ARC NTD contains long helices, predicted to form an anti-parallel coiled coil; binding of ARC to phospholipid membranes requires the NTD. Our data support a role for the ARC NTD in oligomerization as well as lipid membrane binding. These findings have important implications for the structural organization of ARC in distinct functional modalities, such as postsynaptic signal transduction and virus-like capsid formation.

## Introduction

The activity-regulated cytoskeleton-associated protein (ARC) is required for long-term synaptic plasticity and memory formation. ARC binds to distinct protein partners in postsynaptic neuronal compartments to mediate a broad range of effects. In long-term depression (LTD), ARC interacts with components of the clathrin-mediated endocytosis machinery to promote endocytosis of α-amino-3-hydroxy-5-methyl-4-isoxazolepropionic acid (AMPA) receptors (DaSilva *et al.* 2016). In long-term potentiation (LTP), ARC enables stabilization of nascent F-actin in dendritic spines (Messaoudi *et al.* 2007; Nair *et al.* 2017). ARC also acts in the nucleus to regulate gene transcription underlying homeostatic synaptic plasticity (Korb *et al.* 2013). Genetic variants of the ARC protein complex are associated with human general intelligence and schizophrenia risk (Fernández *et al.* 2017; Myrum *et al.* 2017). Thus, current evidence supports a view of ARC as a functionally versatile hub crucial for synaptic plasticity and aspects of human cognition (Nikolaienko *et al.* 2017b).

Structural predictions and biochemical analyses suggest two folded regions flanking a central linker region in human ARC (Myrum *et al.* 2015). The C-terminal domain (CTD) of ARC contains two homologous domains, which may form a bilobar structure (Zhang *et al.* 2015). These small domains, the N- and C-lobe, show structural homology to the HIV Gag capsid protein CA. The N-lobe was co-crystallized with a peptide from the cytosolic tail of the AMPA receptor-binding protein stargazin, and the same binding pocket may accommodate other ligand peptides (Zhang *et al.* 2015), such as those from guanylate kinase-associated protein (GKAP), Wiskott-Aldrich syndrome protein family member 1 (WAVE1), and the glutamate receptor GluN2A. The interaction between ARC and GKAP is compatible with GKAP-driven activity-dependent reorganization of the postsynaptic density (PSD) structural network (Shin *et al.* 2012). WAVE1, a regulator of actin filament branching in dendritic spines, is a potential mediator of ARC in structural plasticity. Interaction with GluN2A further suggests a connection to N-methyl-D-aspartate (NMDA) receptors. The structure and function of the ARC N-terminal domain (NTD) remain unknown thus far.

Full-length recombinant ARC is capable of reversible self-oligomerization in water, while in salt-containing buffers, it is prone to aggregation (Byers *et al.* 2015; Myrum *et al.* 2015). Recently, ARC was shown to form virus-like capsid structures; these capsids contain RNA and are released from neurons in extracellular vesicles (Ashley *et al.* 2018; Pastuzyn *et al.* 2018). Alternating oligomeric states may have functional relevance in ARC signalling and capsid formation, but the details of such mechanisms at the molecular level are lacking. Pure monomeric forms of full-length ARC have not been reported, while being indispensable for detailed structure-function studies.

We purified monomeric full-length human ARC and carried out structural and biophysical characterization using small-angle X-ray scattering (SAXS) and synchrotron radiation circular dichroism spectroscopy (SRCD). Analysis of full-length and truncated forms of ARC revealed a compact structure, in which the oppositely charged N- and C-terminal domains are juxtaposed. Peptide ligands bound specifically to the ARC N-lobe and did not induce structural changes within the lobes or flanking unfolded regions. Furthermore, the NTD of ARC contains two long positively charged helices and mediates binding to lipid membranes, possibly by interaction with phospholipid headgroups. The structural model was validated through Förster resonance energy transfer (FRET) imaging in hippocampal slices. Based on structural evidence, we posit distinct functional modalities for ARC in postsynaptic signal transduction and virus-like capsid formation.

## Materials and Methods

### Recombinant protein production

Full-length human ARC and six truncated forms (Table 1, Table S1, Figure 1A) were expressed in *Escherichia coli* BL21(DE3) cells, resulting in an N-terminally His-tagged protein cleavable by TEV protease. The constructs ARC_206-396_, ARC_207-277_, and ARC_277-370_ were expressed with a maltose binding protein and ARC_170-396_ with a ZZ-domain tag, both located after the His-tag and removed during the purification.

**Table 1.** ARC constructs. The numbering is based on the human ARC sequence. Biophysical parameters obtained using SAXS, SEC-MALS, and DLS for the different recombinant ARC constructs. R_g_* is the R_g_ calculated for a perfect sphere with the *ab initio* volume of the protein.

| Protein name | Description | $R_g$ (nm) | $R_h$ (nm) | $R_g/R_h$ | $R_g/R_g^*$ | $D_{max}$ (nm) | $MW_{calc}$ (kDa) | $MW_{MALS}$ (kDa) | *ab initio* volume/$MW_{MALS}$ (nm<sup>3</sup>/kDa) |
| --- | --- | --- | --- | --- | --- | --- | --- | --- | --- |
| ARC<sub>1-396</sub> (pH 7) | Full-length ARC | 3.58 | N/A | N/A | 1.43 | 14.37 | 47.9 | 50.1 | 2.78 |
| ARC<sub>1-396</sub> (pH 11) | Full-length ARC | 3.85 | N/A | N/A | 1.51 | 15.0 | 47.9 | 52.8 | 2.86 |
| ARC<sub>131-396</sub> | Linker region and CTD | 3.71 | 3.69 | 1.00 | 1.98 | 14.01 | 30.2 | 34.4 | 1.72 |
| ARC<sub>170-396</sub> | Half linker and CTD | 3.03 | 3.25 | 0.93 | 1.60 | 12.20 | 26.4 | 27.8 | 2.17 |
| ARC<sub>206-396</sub> | C-terminal domain | 2.80 | 2.92 | 0.96 | 1.66 | 11.94 | 22.6 | 25.7 | 1.69 |
| ARC<sub>206-361</sub> | CTD without C-tail | 2.25 | 2.40 | 0.94 | 1.41 | 8.74 | 18.7 | 20.7 | 1.79 |
| ARC<sub>207-277</sub> | N-lobe of CTD | 1.43 | 1.92 | 0.74 | 1.24 | 6.13 | 8.9 | 9.7 | 1.37 |
| ARC<sub>277-370</sub> | C-lobe of CTD | 1.69 | 2.03 | 0.83 | 1.21 | 5.95 | 11.3 | 12.3 | 2.00 |

**Figure 1.**
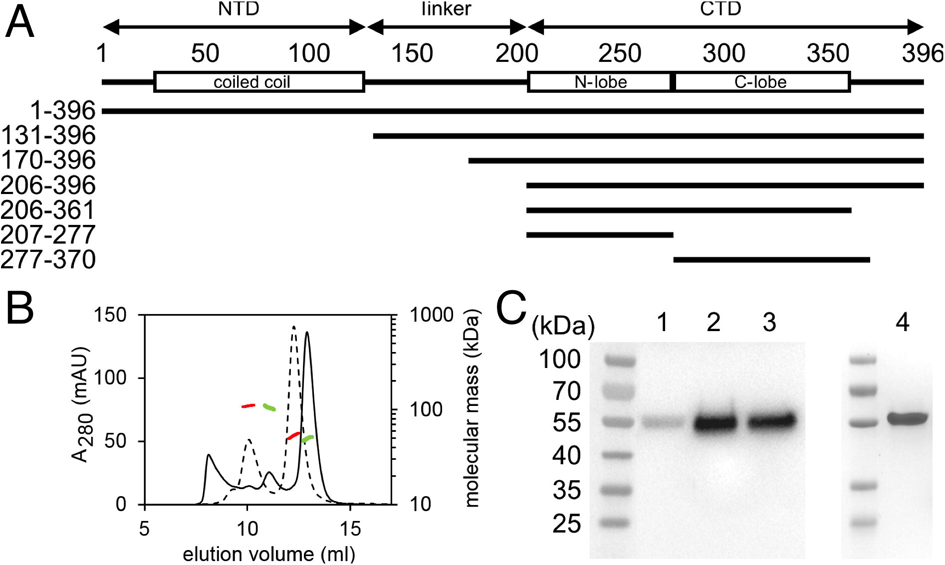
Purification of soluble monomeric ARC. A. The recombinant ARC variants analyzed in this work. B. SEC-MALS analysis of purified full-length ARC at pH 7 (solid chromatogram, mass shown in green) and 11 (dashed, mass shown in red). C. Protein extraction from rat hippocampus analyzed on SDS-PAGE and western blotting using an anti-ARC antibody. 1 - The soluble fraction after homogenization at neutral pH; 2 - The insoluble fraction from neutral pH, re-solubilized in pH 12; 3 - The insoluble fraction solubilized in neutral pH with 2% SDS; 4 - Purified recombinant full-length ARC (Coomassie staining).

For the expression of truncated ARC constructs, cells were grown at +37°C until an A_600_ of 0.6 was reached. Protein expression was induced for 2-4 h at +30°C using 1 mM isopropyl β-D-1-thiogalactopyranoside. The cells were lysed in HBS buffer (40 mM HEPES, 100 mM NaCl, pH 7.5) containing 0.1 mg/ml lysozyme, using one freeze-thaw cycle followed by sonication. In the case of ARC_131-396_, which contains one cysteine, 1 mM DTT was present in all of the purification steps until size exclusion chromatography (SEC).

The lysate was centrifuged at 16 000 g for 30 min at +4°C, and the soluble fraction was loaded onto a Ni-NTA resin, which was washed with HBS containing 20 mM imidazole. The bound protein was eluted with HBS containing 300 mM imidazole and cleaved with His-tagged TEV protease (van den Berg *et al.* 2006). The protein solution was dialyzed for 16-20 h at +4°C against a reservoir of 20 mM HEPES (pH 7.5), 100 mM NaCl, and 1 mM DTT. The dialyzed sample was passed through a Ni-NTA resin for removal of the TEV protease and the cleaved His-tag. The final purification step was SEC, using a Superdex S200 16/600 column equilibrated with TBS (20 mM Tris-HCl, pH 7.4, 150 mM NaCl) for all of the ARC truncation constructs except for ARC_277-370_ and ARC_207-277_, which were purified using a Superdex S75 16/600 column. The monomeric protein peak was collected and concentrated using spin concentrators. In the case of ARC_131-396_, 1 mM DTT was added to the solution before concentration.

The purification of full-length ARC (ARC_1-396_) could not be performed using the protocol above, due to the formation of inclusion bodies during expression. The cells were induced as above, but kept at +37°C for the induction period. After lysis as above, with 1 mM DTT present, the insoluble fraction was washed with HBS containing 1 mM DTT and resuspended in 50 mM CAPS, 50 mM Tris, 10 mM DTT (pH 12). The sample was centrifuged at 16 000 g for 10 min at +25°C, and the soluble fraction was diluted 10-fold with HBS. For protein preparation at high pH, the dilution was made with 20 mM CAPS (pH 11) instead of HBS. The diluted protein solution was loaded onto a Ni-NTA resin, washed with HBS containing 1 mM DTT and 20 mM imidazole before elution with HBS containing 1 mM DTT and 300 mM imidazole. The eluted protein was loaded onto a Superdex S200 16/600 column, equilibrated with TBS, and the monomeric protein peak was collected. In the case of protein preparation at high pH, Tris was again replaced with 20 mM CAPS (pH 11). The monomeric fraction was supplemented with 1 mM DTT and concentrated. The His tag was not removed from full-length ARC.

Protein concentrations were determined by absorbance at 280 nm. Protein purity was analyzed with SDS-PAGE, giving a single strong Coomassie-stained band. Protein identity was confirmed using mass spectrometry of trypsin-digested in-gel samples, as described (Raasakka *et al.* 2015).

ARC-interacting peptides (Zhang *et al.* 2015) were synthesized by GenScript (Hong Kong, China; RRID:SCR_002891) with N-terminal acetylation and C-terminal amidation. The peptides were from Stargazin (RIPSYRYR), GKAP (TSPKFRSR), WAVE1 (RTPVFVSP), and GluN2A (RNPLHNED).

### Size exclusion chromatography – multi-angle light scattering (SEC-MALS)

Absolute molecular masses were determined with MALS, using either a DAWN Heleos, or a miniDAWN Treos MALS detector (for ARC_1-396_ and ARC_207-277_). The SEC column used was either a Superdex S200 Increase 3.2/300 or, in the case of ARC_1-396_ and ARC_207-277_, a Superdex S200 Increase 10/300. The SEC-MALS systems were calibrated with ovalbumin, and the running buffer was TBS. Protein concentration was measured with an online refractometer, and the hydrodynamic radius with either an online quasi-elastic light scattering module or, in the case of ARC_207-277_, a Zetasizer Nano (Malvern Instruments, UK).

### Extraction of native ARC from rat brain tissue

One *Cornu Ammonis* region of the rat hippocampus, corresponding to ∼100 mg of tissue, was homogenized in 400 µl of a buffer containing 50 mM Tris-HCl (pH 7.4), 150 mM NaCl, 1 mM EDTA, 1 mM DTT, and 0.1% Triton X-100, using a Potter-Elvehjem homogenizer and divided in two. The lysate was centrifuged at 21 000 g for 15 min at +4°C. The supernatants correspond to the soluble protein fraction, and the pellets were respuspended in 200 µl of either the above buffer containing 2% SDS or a buffer containing 50 mM CAPS (pH 12), 150 mM NaCl, 1 mM EDTA, 1 mM DTT, and 0.1% Triton X-100. The resuspended samples were centrifuged as above, and the supernatants correspond to proteins dissolved by SDS or high pH, respectively. The protein fractions were loaded on SDS-PAGE gels, and immunoblotting was performed using polyclonal rabbit anti-ARC antibodies and HRP-conjugated anti-rabbit secondary antibodies (Synaptic Systems, Germany; RRID:SCR_013612).

### Circular dichroism spectroscopy

Folding of ARC constructs was followed by SRCD experiments. Truncated ARC was diluted into 20 mM phosphate buffer (pH 7.4) with 150 mM NaF and 0.5 mM DTT, while the full-length construct was desalted using a PD-10 column into the same buffer. For the sample at pH 11, phosphate was replaced with 20 mM CAPS (pH 11). Protein concentrations were 6-66 µM. For mixtures of ARC_207-277_ and ligand peptides, ARC_207-277_ was at 67 µM, and the peptides were present either in equimolar amounts or at a 4-fold molar excess.

SRCD spectra were measured between 170 and 280 nm at +10°C, using a 100-µm quartz cuvette, on the AU-CD beamline at the ASTRID2 synchrotron storage ring (ISA, Aarhus, Denmark). Deconvolutions were made with Dichroweb (Whitmore and Wallace 2004) using CDSSTR (Johnson 1999) and SP175 (Lees *et al.* 2006) to estimate secondary structure content.

CD data comparing the ARC CTD at neutral and high pH and the effect of lipids on the full-length ARC protein were collected using a Jasco J-810 Spectropolarimeter and a 1-mm quartz cuvette at +20°C. The protein concentration for ARC CTD was 5 µM in a buffer consisting either of 20 mM phosphate (pH 7.4) or 10 mM CAPS (pH 11). The protein concentration of full length ARC was 1.5 µM in a buffer consisting of Tris-HCl (10 mM, pH 7.4), NaF (75 mM) with and without liposomes DOPC/DOPS (1:1 ratio, 0.5 mM).

### Small-angle X-ray scattering

SAXS data were collected on the BM29 beamline (Pernot *et al.* 2013) of ESRF (Grenoble, France), the B21 beamline at Diamond (Oxfordshire, UK), the P12 beamline (Blanchet *et al.* 2015) at EMBL/DESY (Hamburg, Germany), and the SWING beamline at SOLEIL (Gif-sur-Yvette, France). The data used to generate models in the absence of bound peptide ligands were collected using a SEC-SAXS setup, where SAXS frames are collected as the protein elutes from a SEC column. The columns used were Agilent BioSEC-3, Superdex S200 Increase 3.2/300, or Superdex S200 Increase 10/300, equilibrated with TBS. For full-length ARC at high pH, the latter column was used with 20 mM CAPS (pH 11) instead of Tris-HCl.

SAXS measurements of protein/peptide mixtures (0.3-10 mg/ml in TBS) were made in batch mode with a 4-fold molar excess of peptide. For the ARC_207-277_ and ARC_277-370_ constructs with peptides, the buffer also contained 2 mM DTT and 2% glycerol. All SAXS measurements were done at +10°C.

SAXS data were processed using either ATSAS (Franke *et al.* 2017) (RRID:SCR_015648) or FOXTROT. The collected frames were checked for radiation damage. The samples measured in batch mode were analyzed at different concentrations to avoid intermolecular events. SAXS models were generated using DAMMIN (Svergun 1999), GASBOR (Svergun *et al.* 2001), CORAL (Petoukhov *et al.* 2012), and EOM (Bernadó *et al.* 2007; Tria *et al.* 2015). In the CORAL run, full-length ARC was modelled by simultaneously including the data from all constructs of different length. The molecular mass of samples in batch mode was based on either absolute scale or a bovine serum albumin standard.

### FRET experiments

#### DNA constructs

FRET sensors for *in vivo* FRET imaging were assembled in the pcDNA3.1(+) vector (Thermo Fisher Scientific, RRID:SCR_008452) behind the CMV promoter. The original polylinker sequence between the *Nhe*I and *Xho*I restriction sites was replaced with a custom polylinker containing a Kozak sequence, start and stop codons, as well as additional restriction sites to facilitate subcloning (5’-GCTAGC-ACTAGT-ACC-ATG-ACCGGT-GCGATCGC-A-GGATCC-GCGGCCGC-A-TCCGGA-TTAATTAA-A-TAA-CTCGAG-3’). The constructs encoding mTurquoise2 (pLifeAct-mTurquoise2, Addgene (RRID:SCR_002037) plasmid #36201) (Goedhart *et al.* 2012), YPet (pCEP4YPet-MAMM, Addgene plasmid #14032) (Nguyen and Daugherty 2005), P2A sequence (pAAV-hSyn1-mRuby2-GSG-P2A-GCaMP6s-WPRE-pA, Addgene plasmid #50942) (Rose *et al.* 2016), and rat ARC (pGEX-4T-3-Arc) (Nikolaienko *et al.* 2017a) have been described previously. The constructs encoding EGFP (pEGFP-N1) and mCherry (pmCherry-C1) were from Takara Bio (Mountain View, CA, USA). A positive FRET control plasmid was obtained by introducing mTurquoise2 and YPet (or EGFP and mCherry) between *Age*I/*Asi*SI and *Bsp*EI/*Pac*I restriction sites, respectively. The resulting plasmid was later used to subclone the P2A sequence (negative FRET control), full-length ARC, the NTD of ARC (ARC_1-140_), the ARC central linker (ARC_135-216_), and the CTD of ARC (ARC_208-396_) between *Bam*HI and *Not*I restriction sites.

#### Slice culture preparation and DNA transfection

Transverse hippocampal slice cultures were prepared from Wistar Hannover GALAS outbred rats (https://www.taconic.com/rat-model/wistar-hannover-galas) at P10±1 and maintained for 12–21 days as described (Gogolla *et al.* 2006; Otmakhov *et al.* 2004) before DNA transfection by single-cell electroporation. For transfection by gene gun, slices were prepared from Sprague Dawley rats at P8-10 and maintained for 7-10 days before transfection.

DNA transfection was performed by single-cell electroporation (Haas *et al.* 2001; Haas *et al.* 2002) for ratiometric FRET experiments and by Gene Gun (Helios) for fluorescence lifetime imaging (FLIM)-FRET experiments. Single slices were transferred to a glass-bottom chamber (Leica) on a motorized shifting table (Luigs and Neumann, Germany), and continuously perfused with artificial cerebrospinal fluid (aCSF) (flow rate ∼3 ml/min) of the following composition: 124 mM NaCl, 2.5 mM KCl, 2 mM CaCl_2_, 1.5 mM MgCl_2_, 1.25 mM NaH_2_PO_4_, 26 mM NaHCO_3_, 10 mM *D*-glucose, balanced with Carbogen gas (95% O_2_, 5% CO_2_), pH 7.4 at room temperature (+22 – +24°C). Electrical pulse parameters for single-cell electroporation were pulse-width 1 ms, −10 V, at 200 Hz for 200-500 ms. After transfection, insert membranes (Millicell Cell Culture Insert, RRID:SCR_015799, Merck Millipore) with slices were returned to the CO_2_ incubator and maintained in culture medium until use. For transfection by gene gun, 1.6-µm gold microcarriers (BioRad, RRID:SCR_008426) were coated with plasmid DNA and fired directly into individual wells in 6-well culture dishes using helium gas (O’Brien and Lummis 2006).

#### Imaging and data analysis

Imaging was performed 48 h after electroporation for all constructs except full-length ARC, which was imaged 24-28 h after electroporation. Slices were placed in the imaging chamber, and FRET ratios were determined after 15-20 min, using a Leica SP5 upright confocal microscope (DMI 6000 CS, Leica Microsystems, Germany; RRID:SCR_008960) with a water immersion objective HCX APO L 20× 1.0 W (Multiphoton, Leica Microsystems, Germany). For excitation of the donor fluorophore, mTurquoise_2_ (mT2), an Argon laser at 458 nm was used. Emission light was separated using an internal AOBS (Acusto Optic Beam Splitter) system, where 465-500 nm light was designated donor emission (mT2) and 525-520 nm light acceptor emission. The signal was captured in separate photomultiplier tubes using XYZ mode bidirectional scanning at 700 Hz, with a pinhole diameter of 180 µm for 35-32 focal steps (1.2-1.5 µm). Each focal plane image (512×512 pixels, 0.5 µm pixel size) was averaged (Line average – 3, Frame average – 3, internal settings of LAS AF software, RRID:SCR_013673).

2-photon FLIM-FRET imaging was performed on an Olympus FV1000 upright microscope through a 60X water immersion objective (NA=1) with an SPC-830 (Becker&Hickl) photon counting board for time-correlated single-photon counting. Photons were counted for 10-20 s, depending on expression level, in 64×64 pixels. The donor (EGFP) was excited using a 910-nm 2-photon laser (Spectra-Physics), and emission was captured in an H7422-40 detector (Hamamatsu). Fluorescence lifetime was calculated in SPC-image software (Becker&Hickl) using mono-exponential curve fitting of photon distributions.

All image processing and analysis for ratiometric FRET experiments was performed in Fiji (RRID:SCR_002285) (Schindelin *et al.* 2012). Raw image files (.lif, LAS AF software) were imported using the Bioformats importer plugin (Linkert *et al.* 2010) and split into two color channels (donor, acceptor). Background signal was subtracted from both channels using the rolling ball background subtraction function before the XYZ stacks were projected to two dimensions (XY) using a sum-intensity projection, and a threshold was manually applied to roughly segment the cells. The image calculator was used to divide the acceptor channel by the donor channel, yielding a 32-bit ratio image. Finally, the mean ratio for each soma was calculated. The data are presented as the mean ± SD of the total sample size.

Negative and positive FRET control constructs were used to define the maximum and minimum FRET achievable for this imaging system and FRET pair. The negative control features the donor and acceptor separated by a P2A sequence, which yields cleavage in the polypeptide during translation. This ensures equimolar expression of donor and acceptor freely diffusing in the cell. In this disconnected state, the local fluorophore concentration is too low for detectable FRET to occur, such that any FRET signal detected is a result of donor/acceptor crosstalk (spectral bleed-through, direct excitation of acceptor) and background signal. In the positive control, the donor and acceptor fluorophores are separated by a 10-amino-acid sequence, short enough to ensure constitutive FRET.

The number of cells used was as follows: ratiometric FRET - ARC_1-396_ (50), ARC_1-140_ (20), ARC_135-216_ (170), ARC_206-396_ (3), positive control (21), negative control (17); FLIM-FRET - ARC_1-396_ (41), ARC_1-140_ (12), ARC_135-216_ (48), ARC_206-396_ (59), positive control (22).

### Isothermal titration calorimetry

A MicroCal iTC200 (Malvern, UK) was used to determine the affinities of peptides to the N-lobe (ARC_207-277_). The peptides (2.2 mM) were injected into the protein solution (0.22 mM) by 26 3-µl injections of the peptide solution, with one initial injection of 0.5 µl and a second 0.5 µl injection after 14 injections due to syringe refill. The protein and the peptides were in TBS buffer, and the experiments were done at +25°C. The data were analyzed with MicroCal Origin 7 (RRID:SCR_002815), using a one-site model.

### Liposome co-sedimentation assay

Liposomes consisting of 1,2-dioleoyl-*sn*-glycero-3-phosphocholine (DOPC), 1,2-dioleoyl-*sn*-glycero-3-phospho-(1’-rac-glycerol)(DOPG),1,2-dioleoyl-*sn*-glycero-3-phospho-*L*-serine (DOPS) and 1,2-dioleoyl-*sn*-glycero-3-phosphoethanolamine (DOPE) were prepared in various combinations by mixing the dissolved lipids in chloroform, drying using a freeze-dryer, and resuspending in TBS. After 15-20 h of agitation at +25°C and 7 freeze-thaw cycles, the suspension was passed through a 100-nm filter 11 times, resulting in 100-nm liposomes. Liposome size was confirmed by dynamic light scattering on a Zetasizer Nano (Malvern instruments, UK).

Co-sedimentation of different ARC constructs was performed by mixing liposomes (0.5 mM DOPC/DOPS, 1:1) and protein (1.5 µM of ARC_1-396_ and 1.9 µM of ARC_131-396_ and ARC_206-396_) in TBS. After 1 h at +4°C, the solutions were centrifuged at 170 000 g for 60 min at +4°C. The supernatant was removed and the pellet resuspended in TBS. Controls were made without liposomes, as well as with 130 mM sodium phosphate (pH 7.4) present. Samples were loaded on 4-20% gradient SDS-PAGE gels and Coomassie-stained.

Co-sedimentation of full-length ARC (3.9 µM) with liposomes were also performed using various lipids (0.5 mM), DOPC/DOPG (1:1), DOPC/DOPS (1:1 and 4:1), DOPC/DOPE (1:1) and DOPC alone. The protein/lipid ratio was varied by increasing the concentration of DOPC/DOPS (1:1) and the effect of NaCl concentration was also tested using DOPC/DOPS (1:1, 0.5 mM) liposomes. The solubility of pelleted liposome/protein complexes (DOPC/DOPS, 1:1, 0.5 mM) after co-sedimentation with full-length ARC (3.9 µM) were tested by resuspension in either TBS pH 7.4 or CAPS (50 mM), Tris (50 mM), pH 12 followed by another centrifugation as described above.

### Notes on study design

The study did not involve pre-registration, randomization, or blinding. A 3D structure of monomeric full-length ARC, validated by live-cell FRET imaging, was considered the primary endpoint of the study. Analysis of ligand peptide and lipid membrane binding by ARC were secondary endpoints. Outliers in SAXS data were removed using standard protocols in the field, incorporated into automated data processing pipelines.

## Results

### Purification of monomeric full-length ARC

In order to facilitate structural studies on full-length ARC and its domains, we set up recombinant expression systems for a number of ARC constructs (Figure 1A, Table 1, Table S1). Bacterially expressed full-length ARC was insoluble under a variety of conditions, as reported before (Byers *et al.* 2015; Myrum *et al.* 2015). However, high pH made ARC soluble and suitable for structural analyses. In the presence of salt and reducing agent, a monomeric peak dominated the size exclusion chromatogram (Figure 1B). The oligomeric state was determined with SEC-MALS and SAXS (Table 1, Figure 1B). The monomeric state was preserved, when pH was reduced to a neutral value. If concentrated above ∼1 mg/ml at neutral pH, ARC slowly aggregated, showing different oligomeric states. At high pH, monomeric ARC could be concentrated without aggregation.

All truncated variants lacking the N-terminal domain were soluble, monodisperse, and monomeric (Figure S1, Table 1), and they could be concentrated to >10 mg/ml. All these variants eluted from SEC as a single peak (Figure S1), and the molecular mass across the peak, determined by MALS, corresponded to a monomer (Table 1). Similarly, modelling the truncated constructs from the single peak by SEC-SAXS showed these variants are monomeric (see below). This indicates that the oligomerizing/aggregating property lies within the N-terminal domain. The NTD alone was insoluble and could not be studied in isolation.

To compare recombinant ARC to ARC *in vivo*, we extracted ARC from brain tissue using similar procedures. Native ARC from rat brain could, indeed, be solubilized at high pH, at levels comparable to SDS extraction (Figure 1C). Tissue lysis at neutral pH solubilized only a minor fraction of ARC. This behaviour of native ARC suggests that the protein is bound to an insoluble component – possibly a membrane - in the cell, through an interaction sensitive to pH and likely involving the N-terminal domain.

### ARC has a helical N-terminal domain sensitive to pH

High-quality SRCD spectra show that full-length ARC contains mainly helical structure (Figure 2A), and deconvolution of spectra from all truncated constructs, as well as selected difference spectra (Figure 2B), allows a detailed mapping of structured regions (Figure 2C). Secondary structure predictions suggest the NTD is helical, while the linker region and C-terminal tail are predicted to be random coils. The CTD and the individual lobe structures give helical spectra (Figure 2A), as expected from crystal structures of the two lobes (Zhang *et al.* 2015). The crystal structure of the N-lobe consists of 59% helix, 7% strand, and 34% random coil, similar to the secondary structure content observed by SRCD (Figure 2C). The C-terminal tail and the linker region produce difference spectra corresponding to random coil (Figure 2B). The NTD gives a strong helical signal, corresponding to >60% helix. The high 222/208 nm CD signal ratio of the NTD difference CD spectrum (Figure 2B) suggests the presence of longer and more ideal helical structures (Greenfield 2006) than those in the CTD. A high 222/208 nm CD ratio is indicative of coiled-coil structure (Alfadhli *et al.* 2002; Kammerer *et al.* 1998; Steinmetz *et al.* 2007), which the NTD is predicted to contain.

**Figure 2.**
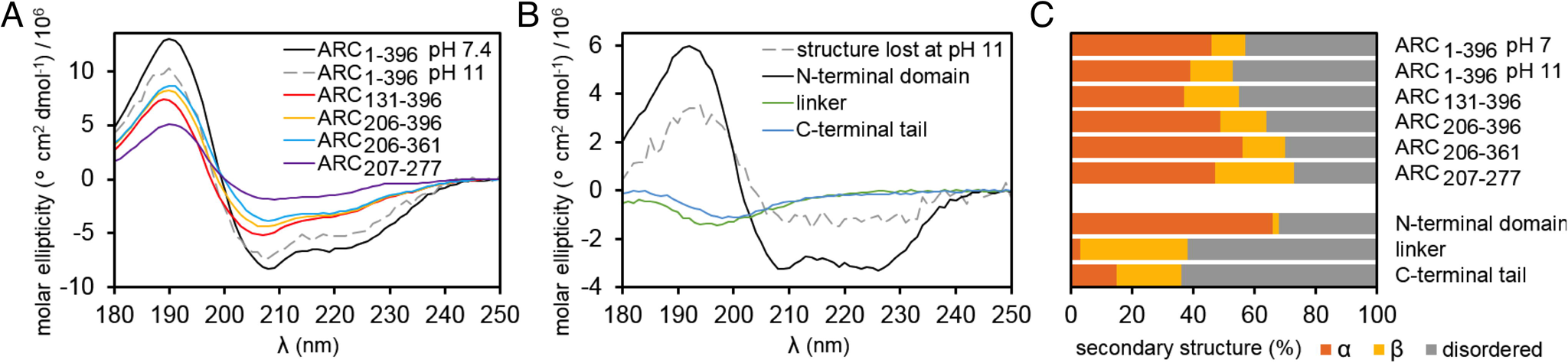
Analysis of ARC folding. A. SRCD analysis of ARC constructs. Note that in all CD spectra, ellipticity has been calculated using the molarity of the protein instead of peptide bonds; this is to facilitate comparison between constructs of different length. B. Difference SRCD spectra. ARC_1-396_ at neutral pH minus the signal at high pH gives the spectrum of the structure that is lost at high pH, ARC_1-396_ at neutral pH minus ARC_131-396_ the spectrum of the N-terminal domain, ARC_131-396_ minus ARC_206-396_ the spectrum of the linker region, and ARC_206-396_ minus ARC_206-361_ the spectrum corresponding to the C-terminal tail. C. Deconvoluted secondary structure contents for all constructs, as well as the segments deduced from difference spectra.

Structural differences between full-length ARC at neutral and high pH were estimated through SRCD difference spectra, showing that most of the secondary structure of ARC remains intact at high pH (Figure 2B,C). The lost helical structure is located in the NTD, as the CTD gave identical spectra at neutral pH and at pH 11 (Figure S2). Thus, a small fraction of the NTD helical structure unfolds at high pH, while ARC remains overall folded. The position of the monomeric peak of full length ARC in SEC shifted depending on pH, appearing slightly larger at high pH (Figure 1B). This increase in the hydrodynamic radius is in line with subtle unfolding. The process is reversible, as the sample at neutral pH was originally solubilized under alkaline conditions.

The 3D structure of the ARC NTD is unknown, and its sequence homology to known protein structures is very low. Secondary structure predictions suggest the presence of two long helices, possibly forming an antiparallel coiled coil (Figure S3). *De novo* fold predictions were made with Robetta (Kim *et al.* 2004). This method gave models with antiparallel, elongated helices (Figure S3). Delta-Blast (Boratyn *et al.* 2012) suggests similarity to the FH2 domain of mDia1 (Nezami *et al.* 2010). This structure consists of two elongated helices. With RaptorX (Källberg *et al.* 2012), the best scoring match is to an SPX domain (Wild *et al.* 2016), again suggesting an elongated structure of two antiparallel helices (Figure S3). The NTD of ARC consisting of elongated helices is in good agreement with the results from both SRCD and SAXS (see below).

### Solution structure of full-length ARC

The individual N- and C-lobes show SAXS profiles expected for globular domains (Figure 3A). For the bilobar structure of the CTD, a dumbbell-like shape is evident; the N- and C-lobes are globular but separated (Figure 3A,B). Full-length ARC has a compact, elongated shape with no signs of isolated globular domains. The maximum diameter of full-length ARC is similar to that of the C-terminal domain (Figure 3B, Table 1), showing that the NTD is located near the bilobar structure of the CTD.

**Figure 3.**
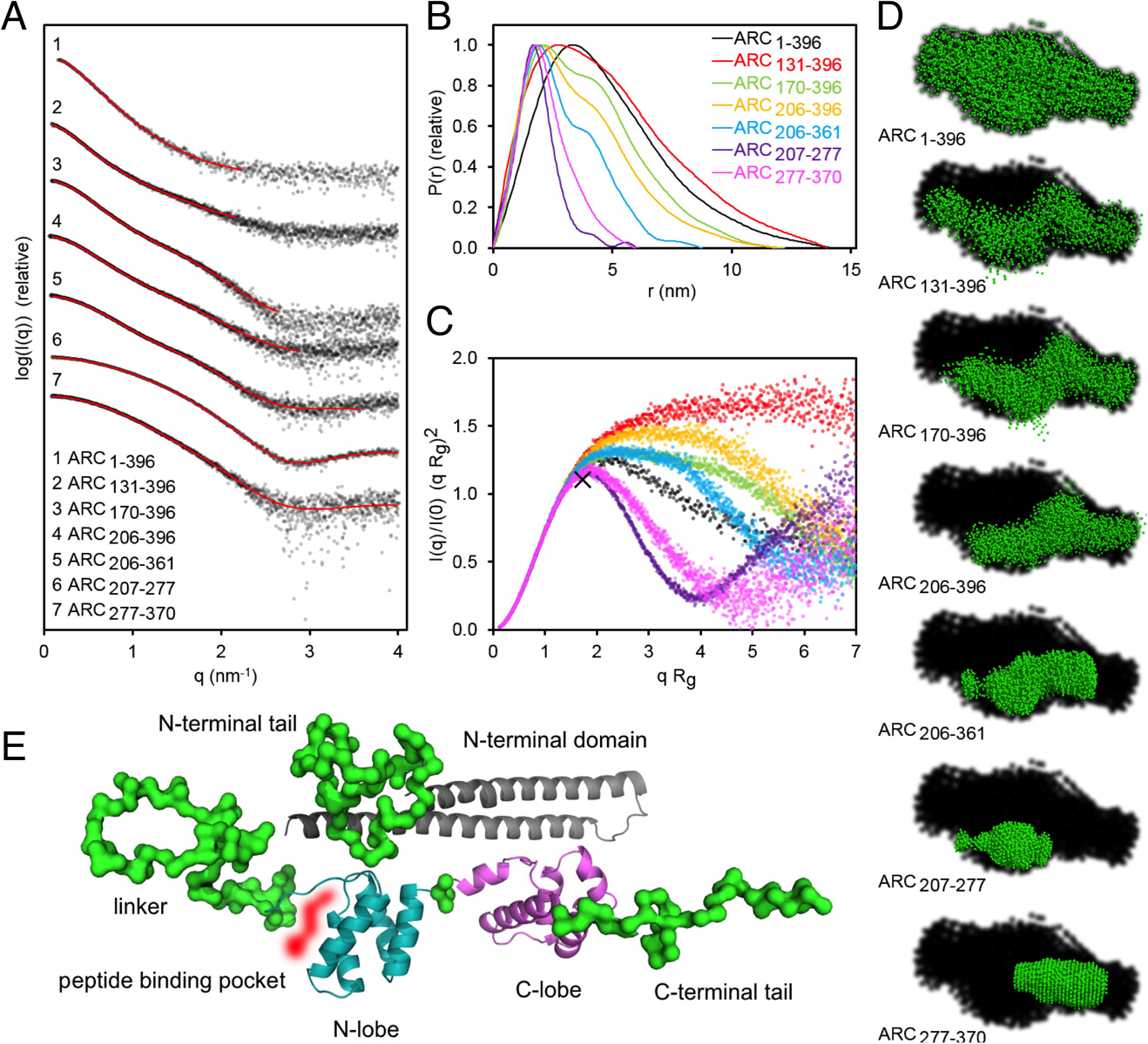
Structural information from SAXS. A. Scattering profiles. The data were shifted on the y axis for better visualization. Fits of *ab initio* models (D) to the data are shown with red lines. B. Distance distribution functions. C. Dimensionless Kratky plots. The cross marks the position of the maximum expected for a rigid, fully spherical particle. D. Dummy atom SAXS models (10 superimposed models per construct) shown in green on a black background, which represents the shape of full-length ARC (ARC_1-396_ χ^2^ =0.9, NSD = 0.6; ARC_131-396_ χ ^2^ =1.3, NSD = 0.8; ARC_170-396_ χ ^2^ =1.4, NSD = 0.7; ARC_206-396_ χ ^2^ =1.4, NSD =0.6; ARC_206-361_ χ^2^ =1.2, NSD = 0.5; ARC_207-277_ χ^2^ =1.4, NSD = 0.5; ARC_277-370_ χ ^2^ =1.3, NSD= 0.4). E. A hybrid model based on SAXS data from all analyzed ARC constructs, the crystal structures of the two lobes, and a homology model of the NTD.

Dimensionless Kratky plots and distance distributions show a single maximum for the two lobes alone, as expected for a well-folded single-domain protein, and the peaks are located near the theoretical position for compact globular particles (Figure 3B,C). The CTD constructs containing both lobes in tandem (ARC_206-361_, ARC_206-396_, ARC_170-393_, ARC_131-396_) show shoulders in such plots, typical for multidomain proteins. Kratky plots indicate a flexible C-terminal tail, and flexibility is increased in constructs with longer central linker regions (Figure 3C). The Kratky plot for full-length ARC, on the other hand, indicates a high degree of folding, compact structure, and little overall flexibility. Thus, by interacting with the CTD, the NTD limits the flexibility of the linker region in the context of full-length ARC. Furthermore, the R_g_ of full-length ARC is slightly smaller than that of ARC_131-396_ (Table 1), and the distance distribution plot shows less tailing for ARC_1-396_ compared to ARC_131-396_ (Figure 3B). Thus, the linker region is stabilized by interactions between the N- and C-terminal domains in full-length ARC. A closed state with the NTD located close to the CTD reduces movements of the linker segment, resulting in a smaller particle diameter of full-length ARC compared to the construct without the NTD.

The available structural data provide no evidence for an open conformation of ARC, whereby the N- and C-terminal domains would be dissociated and connected by an extended linker. This theoretical open state would result in a much larger diameter. As the concentration of full-length ARC is increased and the protein starts to form aggregates, an apparent dimer peak appears in SEC (Figure 1B); this peak could in theory correspond to an open state with a larger R_h_. However, SEC-MALS clearly shows that this peak is a dimer and not an open monomer (Figure 1B). A dimer of ARC could be a relevant first step towards the formation of capsid-like structures.

Various approaches were used to model the 3D structure of ARC based on SAXS data, including dummy atom and chain-like *ab initio* modelling, hybrid modelling coupled with loop building, and prediction of conformational ensembles. Dummy atom models for all constructs were first assembled. The models of the different truncations of ARC can be combined to build a full-length ARC structure (Figure 3D). Full-length ARC is elongated but compact, and the N- and C-terminal halves are located close together. A comparison of full-length ARC with the model of ARC_131-396_, in which the NTD is missing, shows an empty space along the elongated side of the full-length model. This volume corresponds to the predicted size and shape of the NTD.

Further truncations of the linker region and the C-terminal tail remove elongations on both sides of the models, leaving only the two lobes as a dumbbell shape, demonstrating that the lobe structures are two separated domains (Figure 3D). Hybrid models of constructs corresponding to the CTD of ARC with and without the linker region and/or the C-terminal tail (Figure S4) show elongated flexible regions corresponding to the linker and C-terminal tail, in agreement with the SRCD difference spectra for these parts of the protein (Figure 2B).

On one side of the dumbbell-like SAXS model corresponding to ARC_206-361_, an extended feature can be seen, which is also present in the model of the N-lobe (ARC_207-277_) (Figure 3D). This extension could correspond to the more flexible N-terminal part of the N-lobe, supposed to bind peptides by folding onto them (Zhang *et al.* 2015). Chain-like models (Figure S4B) generally agreed with the dummy residue models, apart from the model of the N-lobe, which gave a bad fit at low q (Figure S3C). EOM analysis (Figure S4C,D) of the SAXS data of ARC_207-277_ gives a better fit, indicating a subpopulation of large particles of roughly twice the diameter. Importantly, this extension corresponds to the N-terminal part of the N-lobe that folds on top of the stargazin peptide in the crystal structure (Zhang *et al.* 2015); in the unbound state, it may be flexible and extend away from the globular domain (Figure S4D,E).

A hybrid model of full-length ARC was generated (Figure 3E), based on SAXS data from all constructs, and using the crystal structures of the N- and C-lobe together with the predicted helical structure of the N-terminal domain simultaneously for modelling. This model positions all different domains of ARC, in good agreement with the *ab initio* models. In the model, the coiled coil of the NTD lies on top of the bilobar CTD; the N and C termini of ARC lie at opposite ends. The N and C termini and the linker are likely to be flexible chains, but the central linker is restricted by the arrangement of the folded NTD and CTD.

For understanding the molecular properties of different ARC constructs, the R_g_ values from SAXS, the R_h_ values from dynamic light scattering (DLS), R_g_ values corresponding to a sphere with the observed molecular volume from SAXS (R_g_*), and the molecular volume/mass ratio were compared (Table 1). For a globular particle, one would expect ratios of R_g_/R_h_ = 0.775, R_g_/R_g_* = 1.0, and volume/mass = 1.212 nm^3^/kDa (Erickson 2009). From SAXS experiments, a volume/mass ratio of 1.7 nm^3^/kDa for proteins has been empirically obtained (Petoukhov *et al.* 2012), possibly reflecting the level of hydration and a typical amount of flexibility in a globular protein. Deviations from theoretical values are signs of non-globularity, and flexible regions take up a proportionally large volume. An analysis of these parameters further confirms that the two lobes are globular, while the addition of the linker and C-terminal tail results in non-globular structures. Full-length ARC is more globular than the truncated constructs with disordered regions, but its volume/mass ratio suggests that the flexible regions, indeed, do take up a large volume in solution also in the context of the full-length protein.

Taken together, experimentally derived 3D models of full-length monomeric ARC allow relative positioning of all domains of ARC, showing that the N- and C-terminal halves of the protein are both elongated and interact with each other along the long axis of full-length ARC. Moreover, full-length ARC is less flexible than expected, and the data provide no evidence for an open conformation.

### Corroboration of the structural model of ARC in live neurons

In order to validate the structural model of full-length ARC, we carried out FRET experiments in live neurons with constructs carrying donor and acceptor fluorophores at the N and C termini, respectively (Figure 4,S5). FRET efficiency reflects the distance between the pair of fluorophores, provided that the relative orientation of the fluorophores is constant and the distance is less than ∼10 nm, which is close to the maximum dimension determined for full-length ARC by SAXS (Table 1).

**Figure 4.**
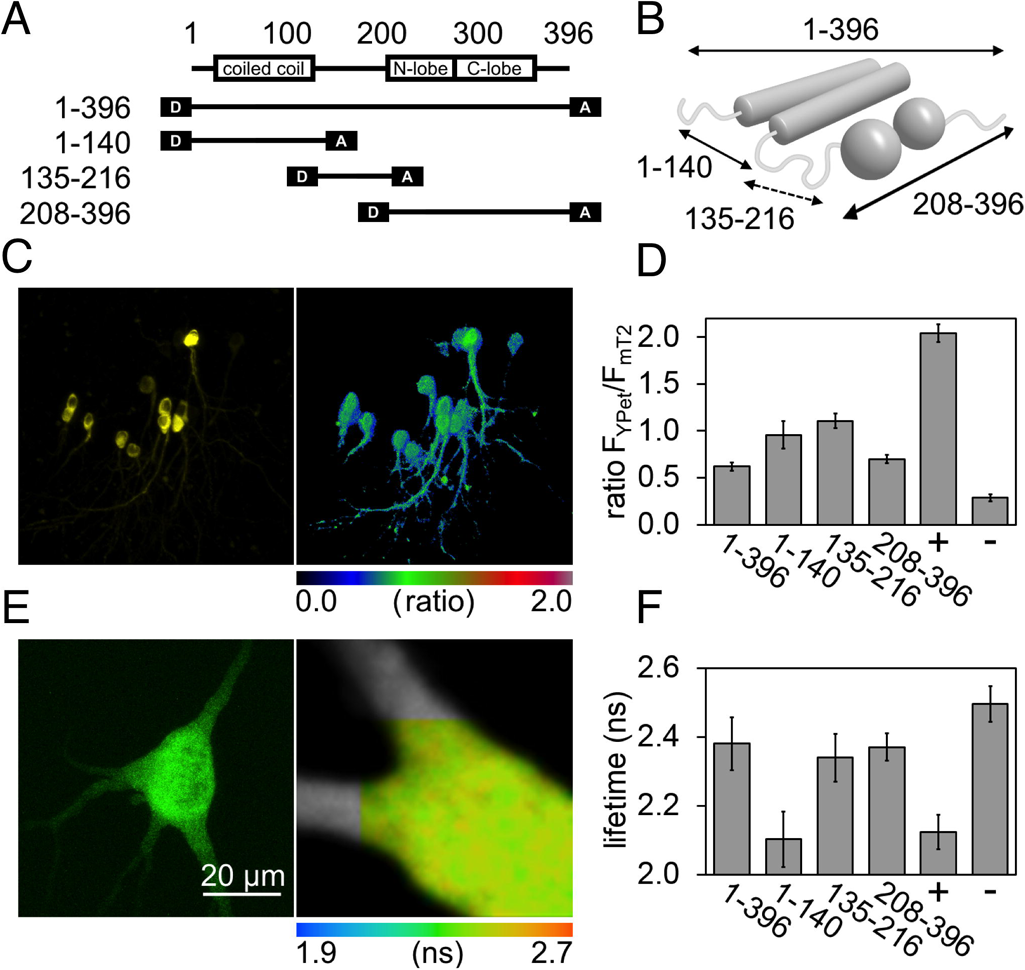
FRET analysis of ARC constructs in hippocampal slices. A. The constructs used in FRET experiments. D, donor; A, acceptor. B. A schematic model of ARC, showing the distances measured with FRET. C. Fluorescence (left) and ratiometric FRET (right) images of neurons expressing full-length ARC. D. Quantification of the ratiometric data (mean ± SD). +, positive control; -, negative control. A high ratio is a sign of a short distance. E. Representative fluorescence (left) and FLIM-FRET (right) images of a neuron expressing full-length ARC. F. Quantitative analysis of FLIM-FRET data. A short lifetime is a sign of a short distance.

Two FRET methods were applied to ARC fragments expressed in cultured rat hippocampal slices (Figure 4A,B). Ratiometric FRET (Figure 4C,D) estimates FRET efficiency non-linearly from fluorescence intensity using an mTurqouise2 (mT2)-YPet pair. FLIM-FRET (Figure 4E,F) provides true FRET efficiency based on the reduced fluorescence lifetime of the donor in the presence of the acceptor, using an EGFP-mCherry (mCh) pair. Both methods show a weak interaction between fluorophores fused to opposite ends of full-length ARC or the CTD (ARC_208-396_), suggesting a distance approaching ∼10 nm. The NTD (ARC_1-140_), on the other hand, shows FRET similar to the positive control, indicating a short distance between the termini. These results corroborate the SAXS-based model of ARC, including a large distance between the N and C termini in full-length ARC and in the CTD, but a short distance between termini of the NTD (Figure 4B). The latter observation fits well with an antiparallel coiled-coil NTD structure formed of two long helices.

The central linker (ARC_135-216_) alone gives rise to high ratiometric FRET efficiency between YPet and mT2, but between EGFP and mCh, as quantified by FLIM, the efficiency is not significantly different from full-length and ARC_208-396_. This disparity is likely caused by the flexible nature of the linker in isolation (Myrum *et al.* 2015), coupled with the ability of YPet to form heterodimers with other members of the GFP family (Kolossov *et al.* 2011; Ohashi *et al.* 2007) - providing, in effect, a measure of flexibility rather than distance. In line with this observation, a recent study combining SAXS and FRET showed decoupling of R_g_ and end-to-end distance in intrinsically disordered molecules (Fuertes *et al.* 2017). Taken together, our FRET experiments validate the experimentally derived 3D molecular structure of full-length ARC.

### Selective ligand peptide binding does not cause large-scale conformational changes

The binding of ligand proteins to the ARC N-lobe in the CTD, next to the linker region, could affect ARC overall conformation and trigger altered functional properties. Peptides known to bind the isolated N-lobe (Zhang *et al.* 2015) (Figure 5A) indeed caused changes in ARC X-ray scattering profiles at low q (Figure 5B), but this could not be linked to clear structural rearrangements (Figure 5C,S6A). When peptides were mixed with ARC constructs and analyzed with SAXS, R_g_ of ARC did not change. However, an increase was seen in forward scattering I(0) for all constructs containing the N-lobe (Figure 5B,C). This increase in molecular mass of the particle is an indication of complex formation between the N-lobe and the peptides.

**Figure 5.**
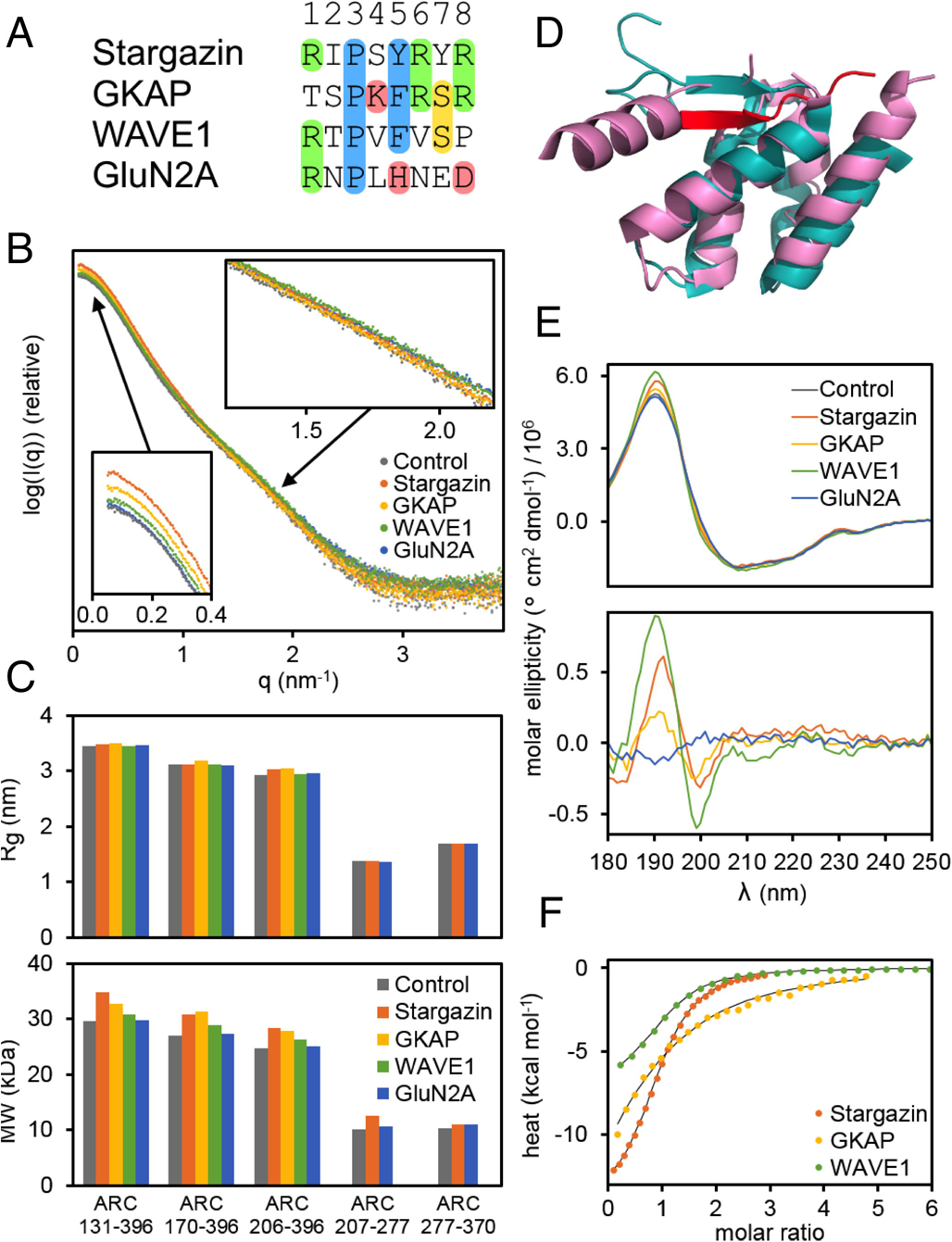
Peptide binding. A. Alignment of the ligand peptide sequences. The consensus motif is coloured in blue, Arg residues in green, and other residues discussed in the text in pink and yellow. B. SAXS scattering curves for ARC_131-396_ with and without peptides. Similar effects were seen with all constructs containing the N-lobe. C. Structural properties of truncated forms of ARC measured with SAXS before and after the addition of peptides. Top, R_g_ obtained through Guinier approximation; bottom, molecular weight of the particle calculated using I(0). D. Superposition of the N- (cyan, PDB entry 4X3H) and C-lobe (pink, PDB entry 4X3X). The stargazin peptide bound to the N-lobe is shown in red. E. SRCD spectra of ARC_207-277_ with and without peptides (top) and difference spectra, in which the spectrum of ARC_207-277_ has been subtracted (bottom). The spectra given by the peptides alone have been subtracted. The peptide:protein molar ratio was 4:1. Plots at a ratio of 1:1 show the same features (Figure S5A). F. Calorimetric titration of the N-lobe with the 3 ligand peptides.

The N- and C-lobe are structurally homologous (Figure 5D); only the N-lobe has been previously tested for peptide binding. The peptide-induced increase in SAXS forward scattering (Figure 5C) occurred for constructs containing the N-lobe (ARC_131-396_, ARC_170-396_, ARC_206-396_, ARC_206-361_ and ARC_207-277_), but not for the C-lobe alone (ARC_277-370_), showing that the peptides specifically interact with the N-lobe. The peptide binding pocket of the N-lobe is not conserved in the C-lobe (Figure 5D), and the function of the structurally conserved C-lobe remains enigmatic.

The increase in I(0) was dependent on the peptide used (Figure 5B,C) and similar responses from all constructs were obtained. Regions outside the ARC N-lobe do not appear to affect peptide binding; this is likely not the case for full-length protein ligands. In SAXS experiments, the stargazin peptide had the strongest effect, followed by GKAP and WAVE1. The GluN2A peptide did not affect forward scattering. These experiments could not be done with full-length ARC, as it requires a SEC-SAXS setup, which is not compatible with rather low-affinity peptide complexes.

SRCD is a sensitive method to detect complex formation even in the absence of apparent secondary structure changes (Cowieson *et al.* 2008). We used SRCD spectroscopy to assess the effect of peptide binding on ARC. After subtraction of the signal of the free peptide, subtle differences can be observed in the spectra (Figure 5E,S6B). Difference spectra (Figure 5E,S6B) show a maximum near 190 nm and a minimum near 200 nm for all peptides except GluN2A, indicative of a similar binding event by the WAVE1, GKAP, and stargazin peptides. In the crystal structure (Zhang *et al.* 2015), the stargazin peptide forms a short β strand between two β strands of the N-lobe (Figure 5D). It is unlikely that the N-lobe can form a stable β sheet in the absence of a bound ligand peptide, and a change in secondary structure should accompany peptide binding. This change should reflect the loss of random coil structure and the gain of β structure. The observed difference spectra reflect overall changes in the protein and peptide components upon complex formation, and while they are reproducible between peptides and experiments, they cannot be used to pinpoint exactly the kind of change occurring.

SAXS and SRCD indicate that the WAVE1, GKAP, and stargazin peptides form a complex with the ARC N-lobe, while the GluN2A peptide does not. ITC was further used to obtain binding affinities (Table 2, Figure 5F). The peptide with the highest affinity was stargazin; binding was also observed for WAVE1 and GKAP. The affinity of the GluN2A peptide was too low to be measured.

**Table 2.** Binding parameters obtained with ITC.

| Peptide | $K_d$ (μM) | $\Delta H$ (kcal/mol) | $-T\Delta S$ (kcal/mol) |
| --- | --- | --- | --- |
| Stargazin | $33.3 \pm 0.8$ | $-14.2 \pm 0.04$ | +8.1 |
| GKAP | $248 \pm 30$ | $-21.4 \pm 0.24$ | +16.4 |
| WAVE1 | $45.7 \pm 1.4$ | $-7.4 \pm 0.04$ | +1.4 |
| GluN2A | N/A | N/A | N/A |

### Binding determinants of ARC ligand peptides

To evaluate differences between ARC ligand peptides, we modelled them based on the stargazin peptide complex (Figure S6C). All peptides can be placed in the binding pocket without severe steric clashes. Most hydrogen bonds between the N-lobe and the stargazin peptide are formed between mainchain groups, indicating that peptide binding may have low sequence specificity. The proposed binding motif is Px(Y/F/H) (Zhang *et al.* 2015), but upon close inspection of the crystal structure, the binding pocket in fact covers 8 residues of the peptide (Figure S6C). The sidechain interactions are clearly related to peptide affinity.

The first residue of the stargazin peptide is an arginine, which is also present in WAVE1 and GluN2A. It is bound between Asp210 and Gln212 (Figure S6C), suggesting that a large polar residue is favoured at this position. The second residue in stargazin is an isoleucine, located in a more hydrophobic environment; the other peptides have small polar residues in this position. Proline at the third position is the only residue present in all four peptides (Figure 5A); it lies in a hydrophobic pocket, and its carbonyl group makes a hydrogen bond to the side chain of His245. This interaction, reminiscent of proteins binding to Pro-rich sequences (Kursula *et al.* 2008), is likely to be a major determinant for peptide binding, and the conserved rigid proline residue defines local backbone conformation in the peptide. The fourth residue in the stargazin peptide is a solvent-exposed serine, unlikely to play a role in specificity. The aromatic residue at position five in stargazin, GKAP, and WAVE1 is in a hydrophobic pocket, making stacking interactions. GluN2A has a histidine at this position, which may explain the observed lack of binding. Arginine residues located at position six and eight of stargazin and GKAP are both near Glu215, allowing salt bridge formation. In the case of GluN2A, this region is instead negatively charged. The stargazin peptide has a tyrosine at position 7, which can stack against Tyr274. These differences explain both the higher affinity of stargazin towards ARC as well as the lack of GluN2A binding.

### The N-terminal domain of ARC binds to phospholipid membranes

ARC may interact with polyanionic surfaces, such as phospholipid membranes, RNA, or microfilaments, especially due to the high positive charge of the NTD. A recent study showed lipid membrane binding and palmitoylation of ARC (Barylko *et al.* 2017). To investigate, which domain of ARC has affinity towards membranes, *in vitro* co-sedimentation assays were performed. Full-length ARC co-sedimented with DOPC:DOPS liposomes (Figure 6A), showing that ARC binds to negatively charged lipid membranes. The NTD was required for ARC binding to liposomes. Co-sedimentation of full-length ARC was prevented by phosphate (Figure 6A), which may compete with protein binding to phospholipids. High salt concentration also prevented co-sedimentation. The results suggest that the interaction between ARC and the membrane is mediated by the positively charged NTD binding to phospholipid headgroups.

**Figure 6.**
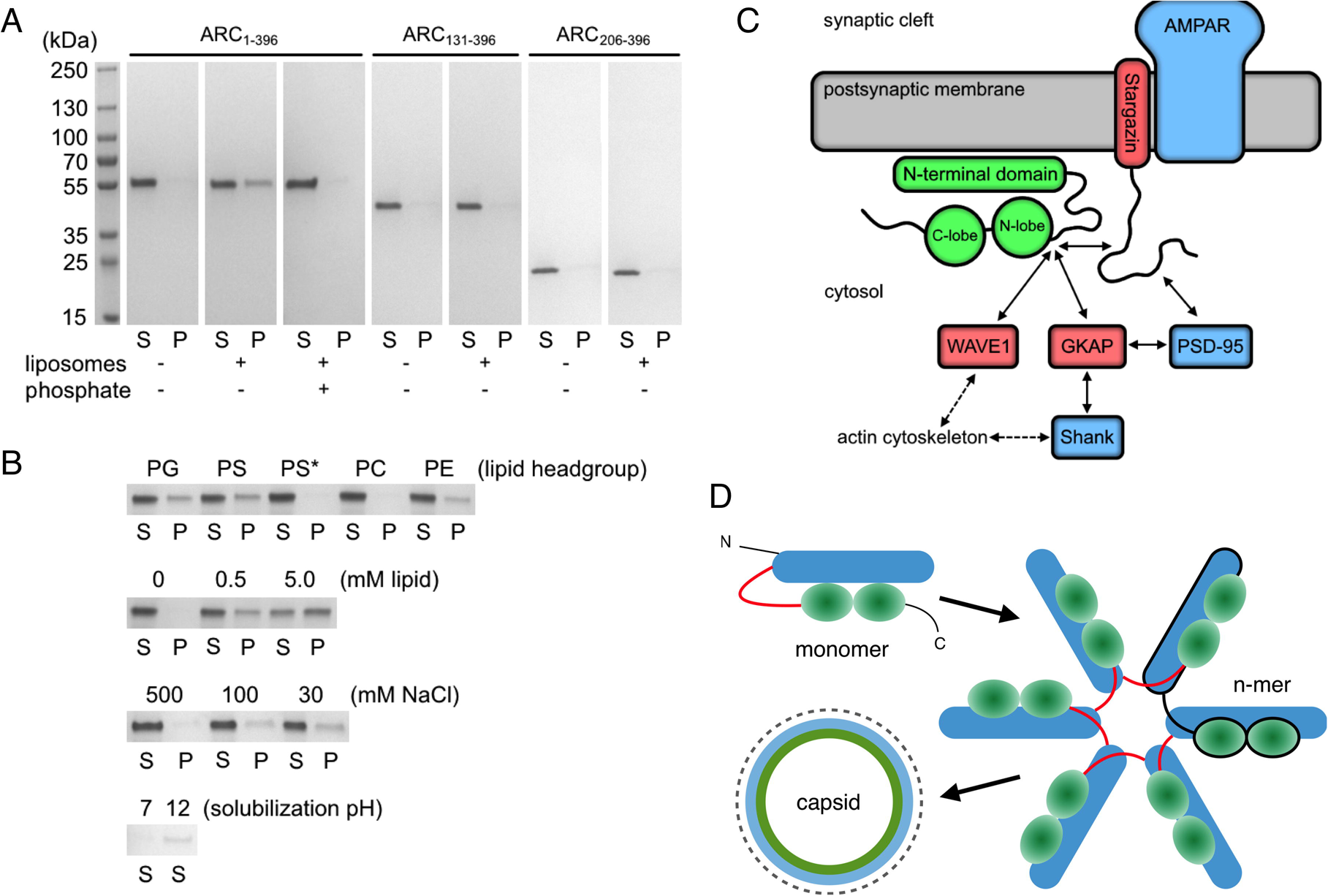
ARC association with lipid membranes. A. Co-sedimentation of ARC with liposomes. B. Effects of lipid headgroup, lipid and salt concentration, and pH on ARC co-sedimentation. C. A model for the localization of ARC. ARC may bind the cytosolic side of the postsynaptic membrane through its NTD; the N-lobe will be in close vicinity to the cytoplasmic tail of stargazin, allowing an indirect connection to the AMPA receptor. Other interactions are likely to compete with this interaction, eventually making ARC a central player in the PSD protein network. D. A possible mechanism of ARC capsid assembly, based on the monomeric full-length ARC structure and analogies to viral capsids. The flexible linker may allow for domain-swapping NTD-CTD interactions between monomers, forming contacts similar to those in monomeric ARC. The symmetry and other details of the ARC assembly in capsid are unknown. The dashed circle around the capsid represents a lipid membrane, and we hypothesize that the ARC NTD forms the outer layer of the capsid under the membrane, while the ARC CTD forms another layer inside the NTD assembly.

The testing of different phospholipid headgroups showed that co-sedimentation did not occur with phosphatidylcholine alone, but the presence of an equimolar amount of DOPS, DOPG, or DOPE promoted binding. Considering the chemical difference between DOPC and DOPE, it thus appears that the methyl groups on phosphatidylcholine, which make its headgroup less polar than that of phosphatidylethanolamine and allows less hydrogen bonding, prevents ARC binding to the membrane surface. To observe pH dependence of ARC-membrane interactions in this system with purified components, the proteolipid pellet was extracted at neutral and high pH. Only high-pH buffer re-solubilized ARC from the lipid pellet, suggesting that, during ARC extraction from recombinant sources or tissue (see above), the membrane interaction must be broken to make ARC soluble. CD spectroscopy indicated that ARC secondary structure content is not affected by the presence of liposomes (Figure S2). Taken together, ARC binds to phospholipid membranes, and this binding is affected by the lipid composition as well as the buffer conditions.

## Discussion

Understanding the molecular function of ARC in different scenarios related to synaptic plasticity, modulation of the protein networks in the PSD, and the formation of virus-like capsids requires detailed knowledge on its 3D structure, interactions, and flexibility. The availability of pure, monomeric, full-length human ARC allows new levels of structural and biophysical characterization.

### Structure of full-length ARC

Full-length ARC is folded and compact, with the helical N-terminal domain located on top of the C-terminal lobes, which are separate entities (Figure 3E). Importantly, no extended linker was observed between the NTD and CTD, and a closed conformation is supported by both X-ray and dynamic light scattering analyses. The N- and C-terminal domains are both elongated, and our data indicate contact between them, although higher-resolution methods will be needed to discern the actual orientations of the domain surfaces with respect to one another.

The structural properties of the model derived from SAXS and SRCD are further supported by intramolecular FRET analysis of ARC expressed in live neurons in cultured hippocampal tissue slices (Figure 4). Both ratiometric and FLIM-FRET data are compatible with a compact arrangement of full-length ARC, in which the CTD and NTD are both elongated. In the isolated CTD, the N- and C-termini lie at opposite ends, whereas the termini of the NTD are in close vicinity, consistent with an antiparallel coiled-coil structure of the NTD. This structural arrangement of the NTD could be critical for interactions of ARC with the polyanionic surfaces of membranes, the actin cytoskeleton, or RNA.

The compact full-length ARC deviates from globularity, as evidenced by the R_g_/R_g_* ratio, and it has significant flexible segments, highlighted by the large molecular volume/mass ratio (Table 1). While the NTD and CTD remain in close contact, the linker and the termini of ARC are flexible, providing an apparent increase in molecular volume in *ab initio* modelling. While ARC has been hypothesized to undergo conformational changes and different states of oligomerization during its functional cycle, it is clear that factors not present in our experiments will be required to trigger conformational changes. Thus, it remains to be seen whether binding events or post-translational modifications can cause opening of the compact structure of ARC.

### Oligomeric status and membrane interactions

ARC has been proposed to alternate between monomeric and oligomeric/aggregated forms (Byers *et al.* 2015; Myrum *et al.* 2015). Recently, ARC was shown to build virus-like capsid structures (Ashley *et al.* 2018; Pastuzyn *et al.* 2018), which are packed into membrane vesicles. Here, we have shown that aggregation tendency and insolubility of recombinant ARC are mediated by the N-terminal domain alone. Upon truncation of this domain, solubility was high, and monodisperse monomers were observed. The C-terminal half of ARC has no propensity to oligomerize on its own. It is possible that the oligomerization property of the ARC NTD plays a role in capsid-like structure formation.

One prominent feature of the NTD is its high isoelectric point of almost 10, something seen for many peripheral membrane proteins (Han *et al.* 2013). Native ARC extracted from brain tissue appears insoluble after homogenization, being assumed to bind to a larger component (Lyford *et al.* 1995). Low solubility may be caused by membrane binding by the N-terminal domain, which could start to aggregate at high protein concentrations. When the NTD forms a helical structure, this will result in an amphipathic arrangement, whereby 24 of the 25 positively charged residues may be located on the same face (Figure S3A); the opposite side would be hydrophobic. The partial disruption or neutralization of this polarized helical structure, *e.g.* by high pH, could be required to solubilize ARC from membrane surfaces.

Phosphate prevented ARC from co-sedimenting with membranes. Interestingly, the SPX domain, a remote homologue of the ARC NTD, functions as a signalling molecule that binds phosphate and inositol polyphophates (Wild *et al.* 2016). Furthermore, capsid formation by ARC was shown to be promoted by phosphate (Pastuzyn *et al.* 2018). Phosphate clearly affects the functional properties of ARC, and likely the positive surface of the NTD plays a role. In addition, co-sedimentation was prevented by high salt concentration, and ARC could be re-solubilized from the lipid pellet at high pH. All the results, including the effects of lipid composition on binding, point towards the importance of polar interactions between ARC and phospholipid headgroups. In a recent paper, recombinant ARC was shown to bind liposomes, and a weak interaction was also observed with phosphatidylcholine alone (Barylko *et al.* 2017). It must be pointed out that the recombinant ARC used in the latter experiments had not been purified by SEC, and therefore, it is likely to have consisted of mostly soluble high-molecular-weight aggregates. In the future, it will be important to determine how phospholipid headgroups affect the formation of ARC-containing endosomes or capsid structures.

The size, shape, membrane binding capability, and presence of long helices in the ARC NTD are all features of BAR domains, which bind to and destabilize the lipid membrane to promote curvature, using positively charged residues (Zimmerberg and McLaughlin 2004). This activity is often involved in endocytosis, and the N-terminal domain of ARC could participate in endocytosis by introducing curvature stress into the membrane surrounding AMPA receptors. In addition, ARC might be involved in recruiting other proteins, such as GKAP and WAVE1, close to membranes (Figure 6B).

### Interaction with ligand proteins

The peptide interaction site (Zhang *et al.* 2015) is located in the globular N-lobe, adjacent to the linker region. Binding to ligand peptides could have structural effects on ARC. From SAXS measurements it was clear that a larger complex was formed after the addition of binding peptides, and a small structural effect could be seen with SRCD. However, SAXS did not reveal restructuring of the linker segment or conformational changes upon peptide binding. When binding a full-length interaction partner, more of the linker region and other domains of ARC are likely to be involved in target protein binding, possibly increasing binding affinity.

Our binding experiments were made with peptides of 8 residues, chosen to match the entire peptide binding site in the crystal structure (Zhang *et al.* 2015). In contrast to stargazin, WAVE1, and GKAP, binding of the GluN2A peptide to the N-lobe could not be observed. This is likely to be caused by poor conservation of the binding motif. The low affinity for this peptide does not rule out an interaction between ARC and GluN2A. The cytoplasmic domain of GluN2A contains several regions with a PxY/F motif, more similar to the stargazin peptide sequence than the sequence suggested earlier to bind ARC (Zhang *et al.* 2015) and analyzed here.

ARC must be located close to the postsynaptic membrane to interact with the short cytoplasmic tail of stargazin. (Figure 6B). Stargazin regulates AMPA receptor trafficking, mobility, and channel properties. Taking into account the binding of multiple stargazin units to the AMPA receptor (Zhao *et al.* 2016), it is possible that ARC surrounds the AMPA receptor, while attached to the membrane through the NTD (Figure 6B). The C terminus of stargazin binds to PDZ domains from several PSD proteins (Chen *et al.* 2000). Therefore, binding of ARC to stargazin might alter stargazin interactions within the PSD scaffold, with impacts on AMPA receptor behaviour. ARC interaction with GKAP is highly relevant in this respect. GKAP is an important hub in the PSD protein scaffold, which will compete with stargazin for ARC binding. GKAP also binds to PSD-95 (Kim *et al.* 1997), and the GKAP C terminus is bound by Shank proteins further down in the PSD assembly (Naisbitt *et al.* 1999; Ponna *et al.* 2018). Thus, ARC binding to the postsynaptic membrane and its ligand proteins may have complex effects on the molecular composition and signalling properties of the PSD.

### Implications of ARC structure for its functional modalities

It is likely that different functions of ARC are related to its various forms and environments, including free monomeric ARC, ARC associated with membranes, endosomes, actin filaments, or nuclear bodies, as well as virus-like ARC capsids. The connections between various ARC binding partners, its post-translational modifications (phosphorylation, SUMOylation, and palmitoylation), membrane binding, and ARC assembly into ordered homo-oligomers, including capsids, are to a large extent still enigmatic.

The high-resolution structure of the ARC-based capsids (Ashley *et al.* 2018; Pastuzyn *et al.* 2018) is not known, including their internal symmetry and details of molecular organization. In the HIV capsid, the Gag capsid protein CA forms both penta- and hexameric assemblies. In both cases, the CA-NTD forms the core of the structure (Pornillos *et al.* 2009; Zhao *et al.* 2013), while the CTD mediates additional contacts to monomeric NTD domains in the CA oligomer (Figure S7). The CA-CTD also keeps the penta- and hexamers together through homodimerization (Zhao *et al.* 2013). It is therefore surprising and noteworthy that the ARC N and C lobes, which are both homologous to the CA-CTD, show no signs of homodimerization. In contrast, we have shown that homo-oligomerization of ARC depends on the N-terminal region of the protein with homology to the Gag MA domain.

The flexible linker between the N- and C-terminal domains in CA is very short. While we observed the ARC NTD and CTD in contact with each other in the case of monomeric ARC in solution, it is perfectly conceivable that the C-terminal lobes of ARC could similarly contact neighbouring ARC monomer NTDs within the ARC capsid structure (Figure 6C). The long flexible linker of ARC would give much conformational freedom in this assembly, possibly allowing domain swapping upon oligomer formation. In line with this hypothesis, the NTD and CTD of ARC were both shown to be required for proper capsid assembly (Pastuzyn *et al.* 2018). We hypothesize that in the ARC capsid-like structures, the NTD forms oligomeric cores linked together through the lobe structures in the CTD. The importance of ARC having two lobe domains homologous to the CA-CTD is currently unknown.

In HIV capsids, the matrix protein (MA) binds to lipid membranes. MA has properties similar to the ARC NTD, including high helical content, positive change, myristoylation, and phosphoinositide as well as membrane binding (Saad *et al.* 2006). The importance of the Arc NTD in membrane binding and oligomeric assembly suggests a dual functional role analogous to both the Gag MA and the CA protein, The interplay between membrane interactions, protein ligand binding to the ARC N-lobe, and capsid formation is an important outstanding issue. It seems unlikely that ligand proteins would specifically get packed into capsids as cargo. However, the presence of ARC ligand proteins and/or a lipid membrane may affect or regulate capsid formation.

## Abbreviations

ARC: activity-regulated cytoskeleton-associated protein
CTD: C-terminal domain
NTD: N-terminal domain
LTD: long-term depression
AMPA: α-amino-3-hydroxy-5-methyl-4-isoxazolepropionic acid
LTP: long-term potentiation
GKAP: guanylate kinase-associated protein
WAVE1: Wiskott-Aldrich syndrome protein family member 1
PSD: postsynaptic density
NMDA: N-methyl-D-aspartate
SAXS: small-angle X-ray scattering
SRCD: synchrotron radiation circular dichroism spectroscopy
FRET: Förster resonance energy transfer
SEC-MALS: size exclusion chromatography – multi-angle light scattering
DOPC: 1,2-dioleoyl-*sn*-glycero-3-phosphocholine
DOPS: 1,2-dioleoyl-*sn*-glycero-3-phospho-*L*- serine
DOPG: 1,2-dioleoyl-*sn*-glycero-3-phospho-(1’-rac-glycerol)
DOPE: 1,2-dioleoyl-*sn*-glycero-3-phosphoethanolamine
DLS: dynamic light scattering
FLIM: fluorescence lifetime imaging
mT2: mTurquoise2
CA: capsid protein
MA: matrix protein
mCh: mCherry

## Acknowledgements

We acknowledge access to and support at SAXS and SRCD beamlines at EMBL/DESY, ESRF, SOLEIL, DIAMOND, and ASTRID2. The project was supported by the Norwegian Research Council (TOPPFORSK grant to C.R.B.).

## Ethics Statement

This research was approved by Norwegian National Research Ethics Committee in compliance with EU Directive 2010/63/EU, ARRIVE guidelines. Persons involved in animal experiments have Federation of Laboratory and Animal Science Associations (FELASA) C course certificates and training.

## Conflicts of interest

The authors declare no conflicts of interest.

## Institutional approval

Institutional approval was not required for this study.

